# Connectomic mapping of pharyngeal and gut sensory circuits in adult *Drosophila*

**DOI:** 10.64898/2025.12.14.694216

**Authors:** Dimitrios S. Giakoumas, Julia M. Zhu, Alaina Jamal, Zepeng Yao

## Abstract

Feeding is regulated by both external sensory signals, such as taste, and internal sensory signals originating from the pharynx and gut. The recent completion of the Full Adult Fly Brain (FAFB) connectome offers an opportunity to map these sensory inputs and their downstream circuits. While the external gustatory receptor neurons have been relatively well characterized, the internal pharyngeal and gut sensory neurons remain less understood. Here, we systemically identify their axonal projections in the FAFB connectome and examine their downstream circuits. We find that the stomodeal nerve, which carries afferent signals from the gastrointestinal tract to the brain, contains multiple types of sensory axons with distinct morphology and downstream output connections. In addition, we identify sensory axons derived from different pharyngeal sense organs and find that chemosensory and mechanosensory axons arborize in distinct regions of the subesophageal zone. Characterization of the second-and third-order neurons reveals the major brain regions that receive input from pharyngeal and gut sensory neurons. Interestingly, a subset of these internal sensory neurons forms monosynaptic connections with various motor neurons and endocrine cells, suggesting that internal signals from the pharynx and gut may directly influence feeding-related motor programs and endocrine output. Together, our study delineates the pharyngeal and gut sensory circuits, laying a foundation for future studies on how internal sensory signals regulate feeding behavior and endocrine functions.

## Introduction

Feeding is one of the most essential behaviors for animals, allowing them to obtain nutrient and energy to support other activities. In many animals, including vertebrate and invertebrate species, feeding is regulated by both external and internal sensory signals. External sensory signals, such as smell and taste, help animals identify potential food sources and evaluate their chemical contents. Taste, or gustation, is particularly important and serves as a checkpoint for feeding ^1–3^. For example, sweet taste indicates the presence of sugars, which are typically nutritive in nature, and promotes feeding. Bitter taste alters the animals of the presence of toxins or harmful compounds, deterring feeding. Once feeding begins, the ingested food continues to be monitored by chemosensory receptors in the pharynx and the rest of the gastrointestinal (GI) tract, among other mechanisms, providing further information about the chemical contents of the food after it is digested ^4,5^. Mechanosensory receptors are also commonly present in the GI tract, providing information about the ingested food volume, among others ^6,7^. These internal GI sensory signals of food content and volume are relayed to the brain via both neural pathways (e.g., the vagus and spinal nerves in mammals) and humoral pathways (e.g., intestinal peptides) to regulate appetite and food intake ^8,9^. While many of the peripheral sensory receptors and pathways have been studied, how central brain circuits process and integrate external and internal sensory signals to regulate feeding is less well understood.

The fruit fly, *Drosophila melanogaster*, has been a valuable model to study the neural regulation of feeding. The gustatory system of flies has been well characterized. Like its mammalian counterpart, it groups the relevant chemicals into a limited number of taste modalities—including sugar/sweet, bitter, water, salt, among others—to guide feeding decisions ^1,10–12^. Gustatory receptors are present in both external and internal sensory organs and tissues, such as the labella and legs (external), and the pharynx and gut (internal), monitoring the chemical contents of the food throughout different feeding stages ^11,13^. While the regulation of feeding by external GRNs has been studied extensively, recent research has demonstrated that internal GRNs located in the pharynx also play critical roles in feeding regulation, detecting a variety of compounds—including sugars, salts, amino acids, and bitter substances—and serving as an additional checkpoint for ingestion ^14–20^. In addition to GRNs, the pharynx also contains mechanosensory neurons, which contribute to the detection of food passage and the regulation of food swallowing ^21–25^. Recent studies have also begun to investigate how gut sensory signals are relayed to the brain to regulate feeding ^26^. One critical pathway involves the enteric stomodeal nerve (StN), which originates from the hypocerebral ganglion (HCG)–corpora cardiaca (CC) region near the anterior portion of the gut ^27–30^. The HCG, for example, contains enteric sensory neurons that detect sugars, sodium salts, and the mechanical distension of the GI tract, and play various roles in feeding regulation ^31–35^. Therefore, *Drosophila* provides an excellent model to study how feeding is regulated by a combination of external and internal sensory signals, a strategy common among many animals.

The recent completion of several connectomes, including the Full Adult Fly Brain (FAFB) connectome for a female fly brain ^36–38^ and two whole-central-nervous-system connectomes for both sexes ^39,40^, offers an exciting opportunity to map the feeding-related sensory neurons and their downstream circuits. Several studies have characterized the external GRNs and investigated their downstream circuitry ^41–47^. The internal sensory neurons from the pharynx and gut, however, remain incompletely mapped in the connectomes. Here, we use the FAFB connectome ^36–38^ to systemically identify internal sensory axons that enter the subesophageal zone (SEZ), a major feeding control center in the fly brain. We identify sensory axons derived from the stomodeal nerve, which relays afferent signals from the GI tract to the brain. We find that these axons consist of multiple subtypes with distinct arborization patterns and postsynaptic partners, suggesting functional specialization. Furthermore, we identify sensory axons originating from different pharyngeal sense organs and find that chemosensory and mechanosensory neurons project to distinct regions of the SEZ. Through analyzing the downstream circuits of the identified gut and pharyngeal sensory neurons, we reveal the top neuropils that receive this sensory information. Interestingly, we discover that some internal sensory neurons form direct synaptic connections onto various motor neurons and endocrine cells, suggesting that their activity may directly influence feeding-related motor programs and endocrine functions. Collectively, our study identifies gut and pharyngeal sensory axons and characterizes their downstream circuits in the fly brain connectome, paving the way for further work on how external and internal sensory signals are integrated to regulate feeding behavior.

## Results

### Identification of putative gut and pharyngeal sensory axons in the FAFB connectome

The subesophageal zone (SEZ) of the *Drosophila* brain receives internal sensory projections from the gastrointestinal (GI) tract and various sense organs located within the pharynx ^29^. The stomodeal nerve (StN), also referred to as the recurrent nerve in some literature, runs along the dorsal surface of the esophagus and contains approximately 43 afferent axons originating from the hypocerebral ganglion (HCG)–corpora cardiaca (CC) region, located near the anterior portion of the gut ^21,28^ (Table 1). (The HCG is also referred to as the stomodeal ganglion in some studies.) After exiting the esophageal foramen, the StN splits into two branches that loop ventrally and fuse with the respective pharyngeal nerves (PhNs) before entering the brain ^27–30^. Because many, but not all, StN axons bifurcate ^28^, slightly fewer than 86 StN axons are expected to enter the brain. Their axonal terminals arborize in the tritocerebrum ^28^, roughly corresponding to the prow (PRW), flange (FLA), and part of the anterior gnathal ganglia (GNG) neuropils defined by Ito et al. ^48^ (Fig. S1).

**Table 1:** Sensory axons in the stomodeal nerve and pharyngeal sensory neurons.

|  | Origin | Composition | Entering brain through |
| --- | --- | --- | --- |
| Stomodeal Nerve (StN) | Gastrointestinal tract | ~43 sensory axons (some bifurcate) <sup>21,28</sup> | PhN <sup>21,28</sup> |
| Dorsal Cibarial Sense Organ (DCSO) | Pharynx | 2 x 6 gustatory neurons <sup>21,22</sup> | PhN <sup>21,29</sup> |
| Ventral Cibarial Sense Organ (VCSO) | Pharynx | 2 x 8 gustatory neurons <sup>21,22</sup> | aPhN <sup>21,29</sup> |
| Labral Sense Organ (LSO) | Pharynx | 2 x 10 gustatory neurons;<br>2 x 8 mechanosensory neurons <sup>21,22</sup> | aPhN <sup>21,29</sup> |
| Fishtrap Bristles | Pharynx | 2 x (~18-23) mechanosensory neurons <sup>21,22</sup> | aPhN <sup>21,29</sup> |

The pharynx contains several sense organs: the dorsal cibarial sense organ (DCSO), the ventral cibarial sense organ (VCSO), the labral sense organ (LSO), and two rows of fishtrap bristles (also called satellite bristles) ^21,22^. The DCSO contains 12 gustatory neurons; the VCSO contains 16 gustatory neurons; the LSO contains a total of 20 gustatory neurons and 16 mechanosensory neurons; and the fishtrap bristles contain around 36 to 46 mechanosensory neurons ^21,22^ (Table 1). DCSO neurons project through the pharyngeal nerves (PhNs) into the brain, while VCSO, LSO, and fishtrap bristle neurons project through the accessory pharyngeal nerves (aPhNs) ^21,29^ (Table 1). The gustatory neurons from the DCSO, VCSO, and LSO project their axons to the dorsal SEZ ^14–16,49–51^, primarily corresponding to the prow (PRW) neuropil (Fig. S1). The mechanosensory neurons from the pharyngeal sense organs are less well characterized, and their axonal projection patterns in the brain remain less clear.

In the FAFB connectome (FlyWire version v783, available from Codex https://codex.flywire.ai), there are 91 sensory axons in the PhNs and 85 sensory axons in the aPhNs ^36–38^ (Fig. 1A-B’’). The aPhN axons are divided into two subclasses in FlyWire Codex: group 1 axons (aPhN1, n = 33) arborize in a more dorsal region of the SEZ and are all annotated as gustatory, while group 2 axons (aPhN2, n = 52) arborize in a more ventral region, and the sensory modality of most neurons in this group is unknown (Fig. 1B-B’’). The PhN axons also contain two groups: 79 axons form one bundle innervating a more anterior region of the SEZ (Fig. 1A-A’’, green), and 12 axons form a separate bundle innervating a more posterior region (Fig. 1A-A’’, red). Interestingly, the bundle of 12 PhN axons is entirely gustatory (similar to aPhN1 axons), and these axons innervate the same dorsal SEZ region (primarily within the prow neuropil) that is also innervated by the gustatory aPhN1 axons (Fig. 1C-C’’). Given that the DCSO contains 12 gustatory neurons projecting through the PhNs and the VCSO and LSO together contain a total of 36 gustatory neurons projecting through the aPhNs (Table 1), and that their axons arborize in the dorsal SEZ ^14–16,49–51^, the 12 PhN axons that form a discrete bundle are likely from the DCSO neurons (Fig. 1A-A’’ and C-C’’, red) and the 33 aPhN1 axons are likely from the VCSO and LSO gustatory neurons (Fig. 1B-C’’, yellow). The slight mismatch in axon numbers for the VCSO and LSO gustatory neurons (36 versus 33) may reflect individual developmental variations or imperfect classification of aPhN axon subclasses in FlyWire. The remaining 79 PhN axons likely originate from the StN, well matching the expected number of StN axons (n < 86) and the brain region they innervate (Fig. 1A-A’’, green). Lastly, given that the pharyngeal sense organs contain around 52 to 62 mechanosensory axons projecting through the aPhNs into the brain from the LSO and fishtrap bristles (Table 1), the 52 aPhN2 axons are likely from these pharyngeal mechanosensory neurons (Fig. 1B-B’’, orange), awaiting confirmation from future studies.

**Figure 1.**
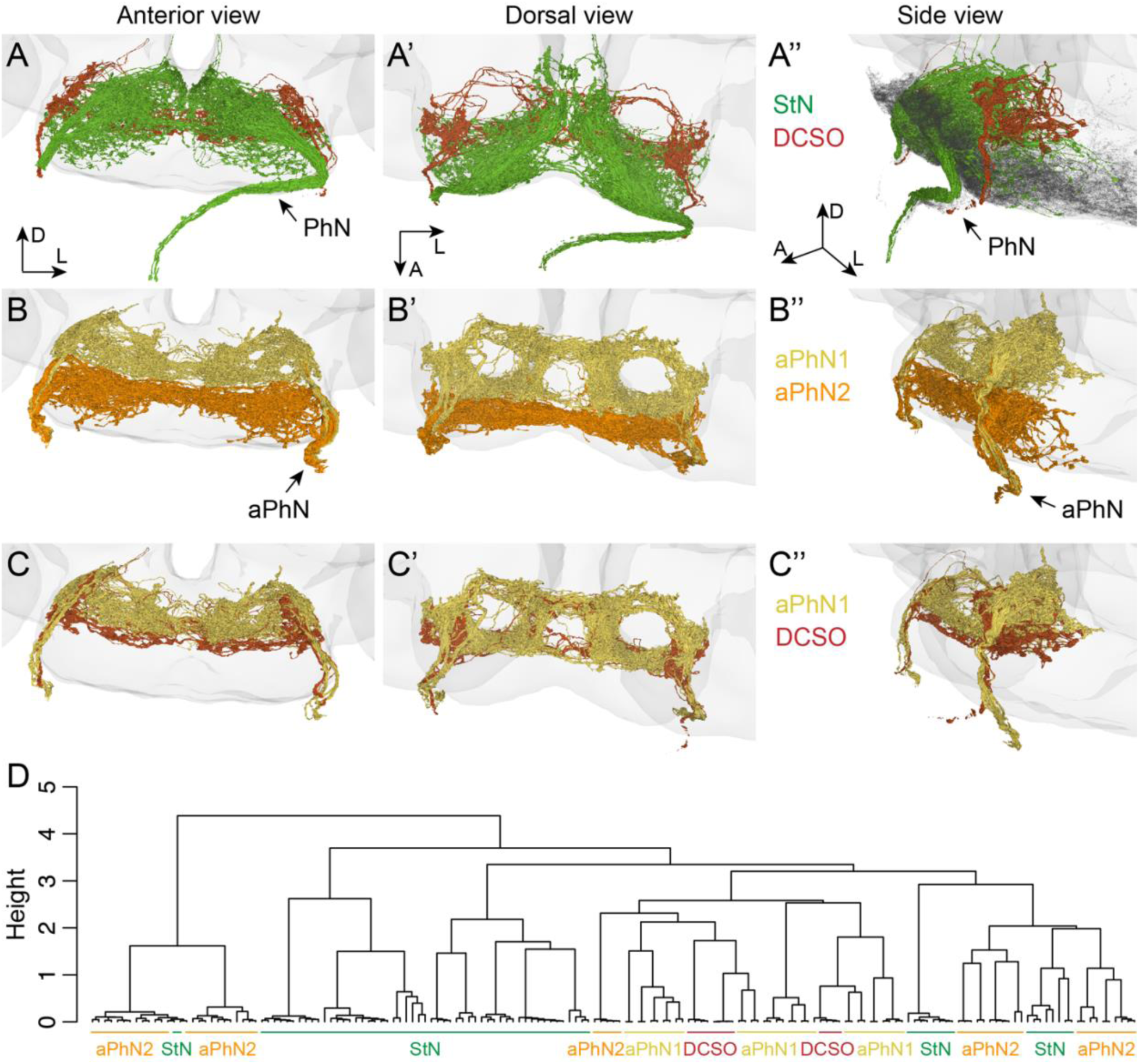
Identification of gut and pharyngeal sensory axons in the FAFB connectome. (**A-C’’**) Anterior views (left panels), dorsal views (middle panels), and side views (right panels) of sensory axons in the pharyngeal nerves (PhN) and accessory pharyngeal nerves (aPhN). (**A-A’’**) show the overlay of StN and DCSO axons; the dark gray shading in (**A’’**) indicates motor neurons in the PhN. (**B-B’’**) show the overlay of aPhN1 and aPhN2 axons. (**C-C’’**) show the overlay of aPhN1 and DCSO axons. In all panels, the light gray shading indicates the brain. The axes indicate the brain orientation: A, anterior; D, dorsal; L, lateral. (**D**) Dendrogram showing the clustering of gut and pharyngeal sensory axons based on the cosine similarity of their postsynaptic partners. The different sensory groups are labeled as indicated.

In addition to anatomical analyses, we investigated the output connections from the PhN and aPhN sensory axons. We retrieved the postsynaptic partners for each PhN or aPhN axon from the connectome and clustered these axons based on the similarity of their output partners using cosine similarity (see Methods for details). The putative DCSO axons cluster tightly with the aPhN1 axons (likely VCSO and LSO gustatory axons) (Fig. 1D), suggesting that these putative pharyngeal gustatory neurons have similar output connections. Interestingly, the putative StN axons cluster interspersed with the aPhN2 axons (likely pharyngeal mechanosensory axons) (Fig. 1D), suggesting that they may converge onto similar downstream circuits.

### Classification of StN sensory axons based on their output synaptic connectivity

The StN axons exhibit different arborization patterns, some diffuse and others more restricted, suggesting the presence of different subtypes ^28^ (Fig. 2). Given that these are sensory axons, we compared their output connections to reveal potential functional subtypes. To this end, we clustered the StN axons based on the similarity of their postsynaptic partners, using cosine similarity (Fig. 2A). Eight StN axons lacking output synapses were excluded from this analysis (Fig. 2B)—this likely reflects a limitation of the automatic synapse detection in the FAFB connectome ^52^ rather than a true absence of output synapses from these sensory axons.

**Figure 2.**
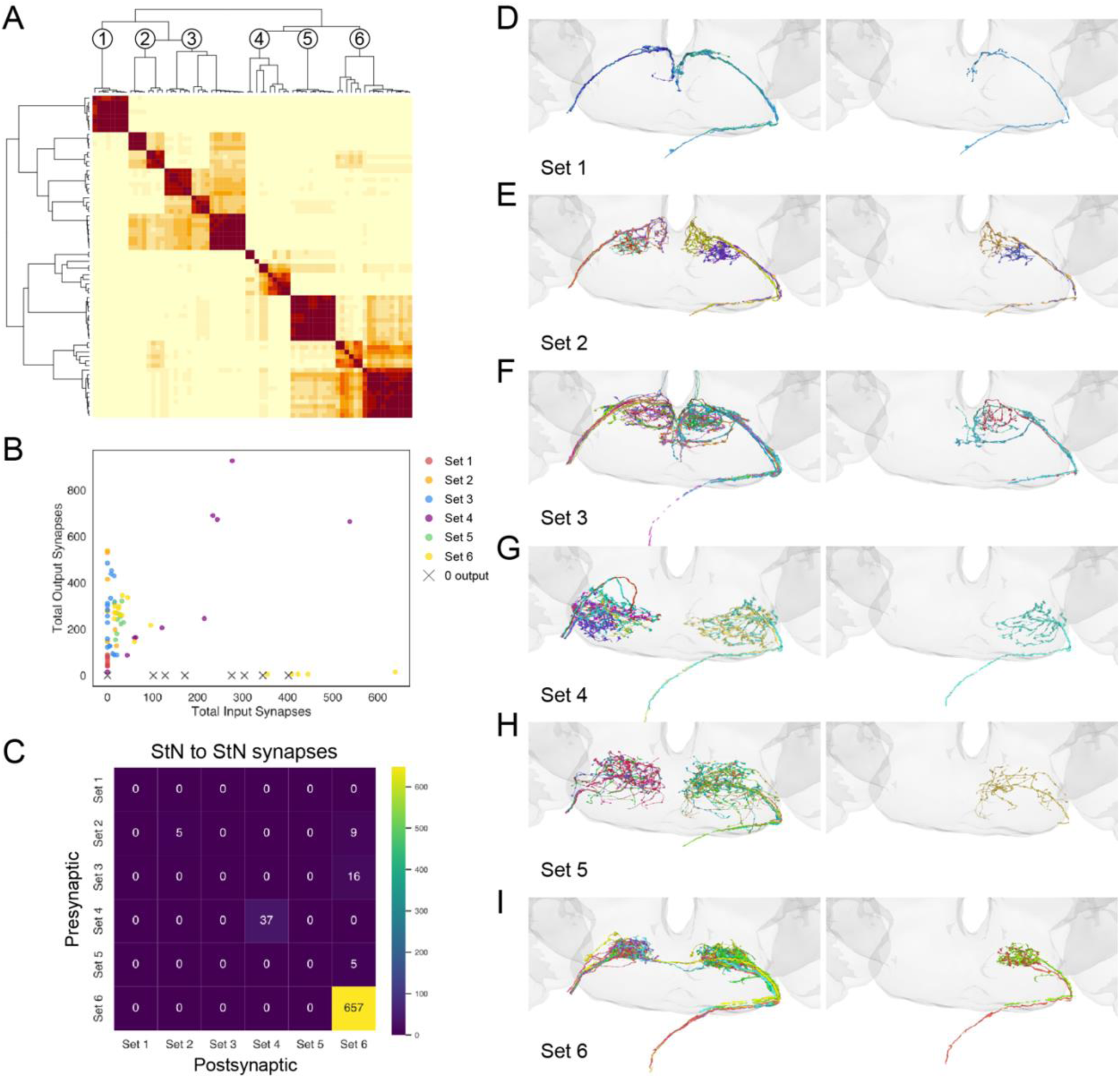
Classification of StN sensory axons based on their output synaptic connectivity. (A) Cosine similarity matrix of StN axons based on the similarity of their postsynaptic partners. Darker colors indicate higher similarity between axons. Six sets are identified, as indicated on the dendrogram. (B) Total output synapses (y-axis) versus input synapses (x-axis) of individual StN axons. The different sets are indicated by different colors. Eight axons lacking predicted output synapses are indicated by crosses (X). (C) Matrix showing the number of synaptic connections between all pairwise sets of StN axons. (**D-I**) Anterior views of different sets of StN axons. Left panels show all the axons; right panels show representative individual axons. For sets 2, 3, and 6, two individual axons are shown to illustrate the presence of distinct morphologies. In all panels, the light gray shading indicates the brain.

Using an intermediate cutoff threshold, our clustering analysis reveals six subtypes of StN axons (sets 1-6) (Fig. 2A). Interestingly, each of these subtypes—identified by functional output connectivity—also appears to be structurally distinct, displaying different axonal terminal patterns and innervating different subregions in the SEZ (Fig. 2D-I). In contrast to the external gustatory receptor neurons (GRNs), whose axonal terminals synapse with each other within the same subtype (e.g., sugar GRNs form synapses with other sugar GRNs) ^41,44^, most StN subtypes have few or no such intra-subtype synapses (Fig. 2C). The exceptions are the Set 6 and Set 4 StN axons, which display high and moderate levels of intra-subtype synapses, respectively (on average, 38.6 synapses/axon for Set 6 and 3.7 synapses/axon for Set 4) (Fig. 2C). Because the StN is poorly characterized, future studies are needed to determine whether these StN subtypes have different molecular identities or physiological roles.

### Downstream circuits of the StN sensory axons

To investigate the downstream circuits of the StN sensory axons, we used the connectome to identify their second-order and third-order neurons. Similar to a recent study on the GRN connectomics ^44^, we define the StN second-order neurons (2Ns) as cells (or root IDs, to be precise) directly postsynaptic to the StN sensory axons, excluding the first-order StN sensory axons. Likewise, the StN third-order neurons (3Ns) are cells (or root IDs) postsynaptic to the StN 2Ns, excluding the first-order StN sensory axons and StN 2Ns. Consistent with many other studies in the field, a minimum of five synapses is used as a threshold for synaptic connections to reduce false positive connections ^37^.

Different StN subtypes have largely distinct 2Ns (Fig. 3A-C), consistent with their classification based on output synaptic connectivity (Fig. 2). For example, the Set 1 StN axons have 13 2Ns, none of which are postsynaptic to the other subtypes of StN axons (Fig. 3B-C), suggesting that they form unique output circuits. The other StN axon subtypes also have a large proportion of 2Ns unique to each subtype (ranging from 42.9% to 83.6%), while other 2Ns are shared among two or more subtypes (Fig. 3B-C). These results suggest that the StN subtypes identified here may represent unique information channels with distinct output circuits, while a small proportion of 2Ns may serve to integrate signals from multiple channels. Most of the StN 2Ns are central neurons confined to the central brain, while other superclasses, such as ascending and descending neurons, are also present (Fig. 3D).

**Figure 3.**
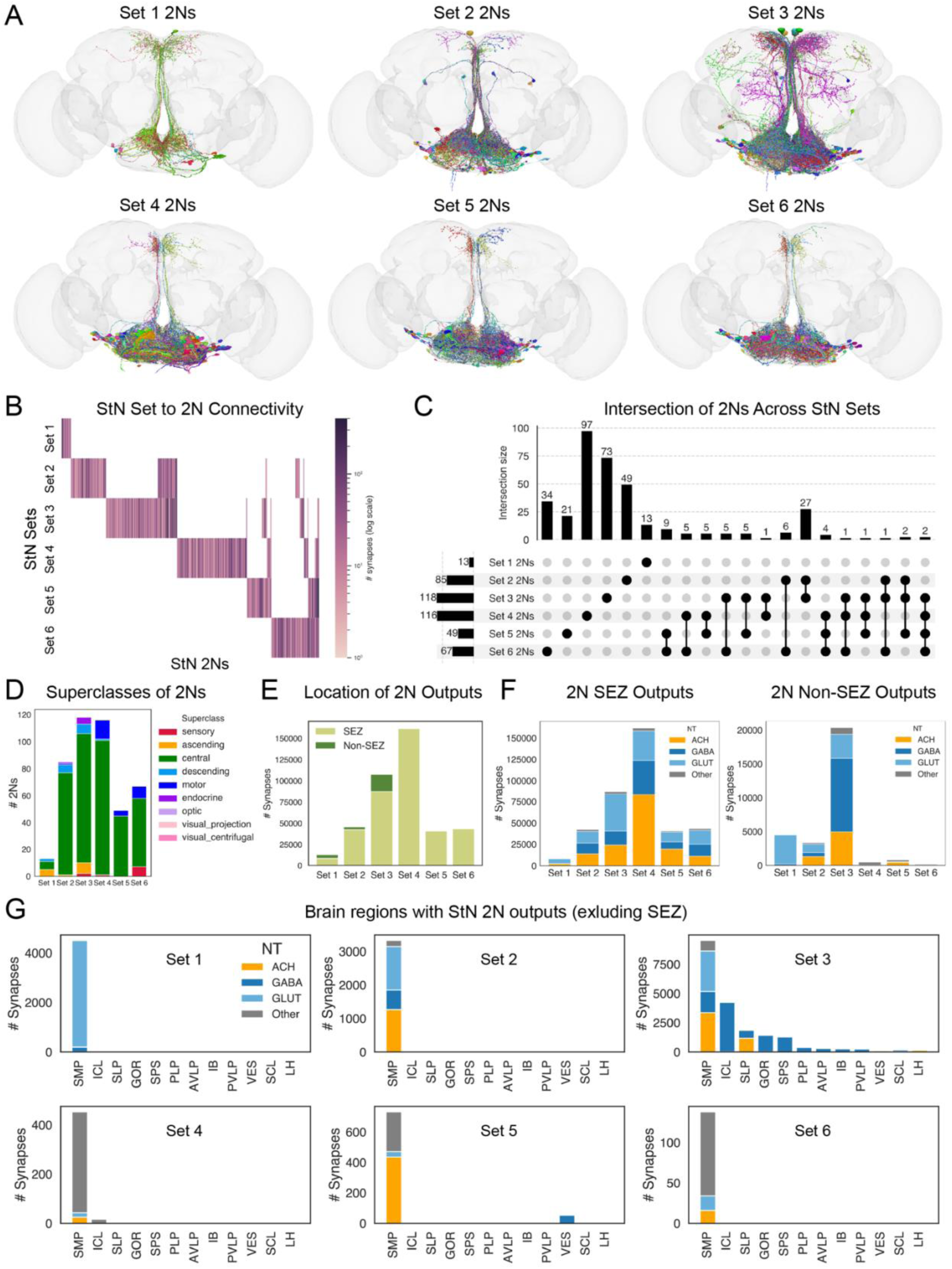
Analyses of StN second-order neurons (2Ns). **(A)** Anterior views of the 2Ns of different sets of StN axons. The light gray shading indicates the brain. (**B-C**) Heatmap (**B**) and UpSet plot (**C**) showing the extent of overlap in 2Ns among different sets of StN axons. The colors in (**B**) are scaled relative to the number of synapses from each StN set to each 2N. (D) The number of 2Ns belonging to different superclasses. (E) The number of 2N output synapses residing within versus outside the SEZ. (F) The predicted neurotransmitters (NT) used by the 2N output synapses that reside within the SEZ (left panel) versus outside the SEZ (right panel). ACH, acetylcholine; GABA, gamma-aminobutyric acid; GLUT, glutamate. (G) The number of 2N output synapses residing in the indicated non-SEZ brain neuropils for each set of StN 2Ns. The predicted neurotransmitters (NT) are indicated by different colors.

Interestingly, some endocrine cells are postsynaptic to Sets 2-3 StN axons, while some motor neurons are postsynaptic to Sets 4-6 StN axons (Fig. 3D). These results indicate the presence of direct sensory-endocrine and sensory-motor connections, which we will explore further in later sections.

The majority of StN 2Ns reside in the SEZ, with most of their output synapses in the SEZ (Fig. 3A and E). The StN 2Ns are predicted to use a variety of neurotransmitters, including acetylcholine (ACh), gamma-aminobutyric acid (GABA), and glutamate (Glut), although these predictions may not always be accurate ^53^ (Fig. 3F). Outside the SEZ, the superior medial protocerebrum (SMP) is the predominant neuropil that receives inputs from all subtypes of StN 2Ns (Figs. 3A, 3G, and S1). While the Set 3 StN 2Ns also target additional neuropils, other StN 2N subtypes almost exclusively target the SMP besides the SEZ (Fig. 3A and G). These results imply the importance of the SMP in processing internal signals from the StN.

At the 3N level, all StN subtypes reach broader networks consisting of hundreds to thousands of neurons (Fig. 4A-B). A relatively small proportion of 3Ns are unique to each subtype (52.6% for Set 3 and 6.0–33.7% for others), while many 3Ns are shared among two or more subtypes (Fig. 4B-C). These results suggest that signal integration across StN subtypes may be widespread at the 3N level. The StN 3Ns consist of different superclasses and utilize different neurotransmitters (Fig. 4D and F). The majority of output synapses from most 3N subtypes are located within the SEZ, except for Sets 1 and 3, approximately half of whose output synapses are located outside the SEZ (Fig. 4E). The Set 3 3Ns, in particular, target a wide range of neuropils (Fig. 4A and G). Outside the SEZ, the SMP remains the predominant brain region that receives synaptic inputs from all StN 3N subtypes (Fig. 4G), further implicating its importance in the StN output circuits.

**Figure 4.**
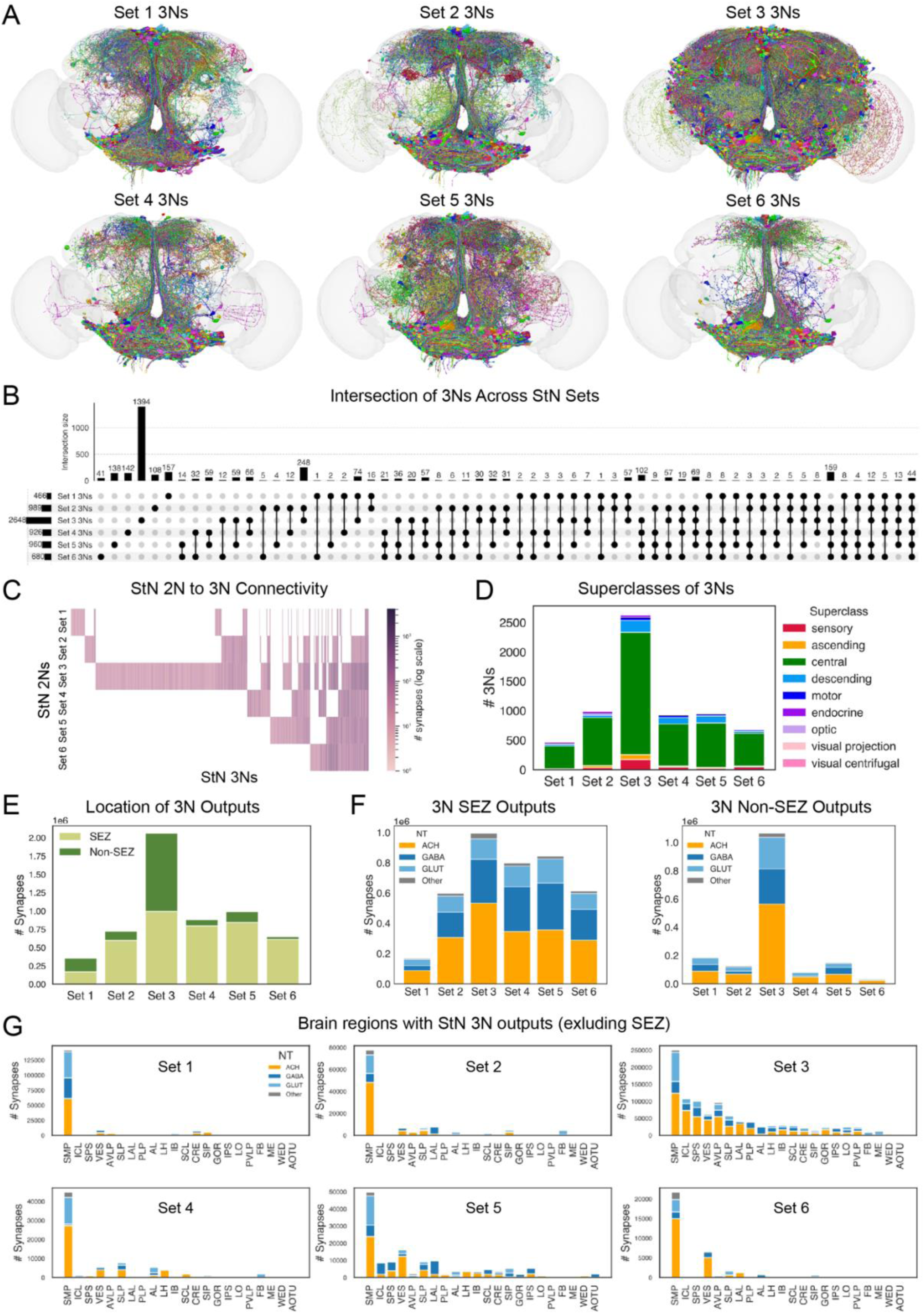
Analyses of StN third-order neurons (3Ns). **(A)** Anterior views of the 3Ns of different sets of StN axons. The light gray shading indicates the brain. (**B-C**) UpSet plot (**B**) and heatmap (**C**) showing the extent of overlap in 3Ns among different sets of StN axons. The colors in (**C**) are scaled relative to the number of synapses from each set of StN 2Ns to each StN 3N. (**D**) The number of 3Ns belonging to different superclasses. (**E**) The number of 3N output synapses residing within versus outside the SEZ. (**F**) The predicted neurotransmitters (NT) used by the 3N output synapses that reside within the SEZ (left panel) versus outside the SEZ (right panel). ACH, acetylcholine; GABA, gamma-aminobutyric acid; GLUT, glutamate. (**G**) The number of 3N output synapses residing in the indicated non-SEZ brain neuropils for each set of StN 3Ns. The predicted neurotransmitters (NT) are indicated by different colors.

### Downstream circuits of the pharyngeal sensory axons

We next investigated the downstream circuits of the sensory axons from the pharynx, the putative DCSO axons, and the aPhN1 and aPhN2 axons (Figs. 1 and 5A). Most of these pharyngeal sensory axons contain both input and output synapses (Fig. 5B), suggesting that they may be modulated by brain inputs in addition to sending out sensory signals. One aPhN2 axon without any output synapses identified was excluded from the subsequent analyses (Fig. 5B). The DCSO and aPhN1 axons, which are likely gustatory axons from the pharynx, form moderate levels of intra-group connections (on average, 5.7 synapses/axon for DCSO-DCSO and 10.6 synapses/axon for aPhN1-aPhN1 connections) (Fig. 5C), comparable to those observed for the external GRNs ^41,44^. In contrast, the aPhN2 axons, which are likely mechanosensory axons from the pharynx, form very few synapses with each other (0.2 synapses/axon on average) (Fig. 5C). These findings imply different strategies for processing gustatory versus mechanosensory signals.

**Figure 5.**
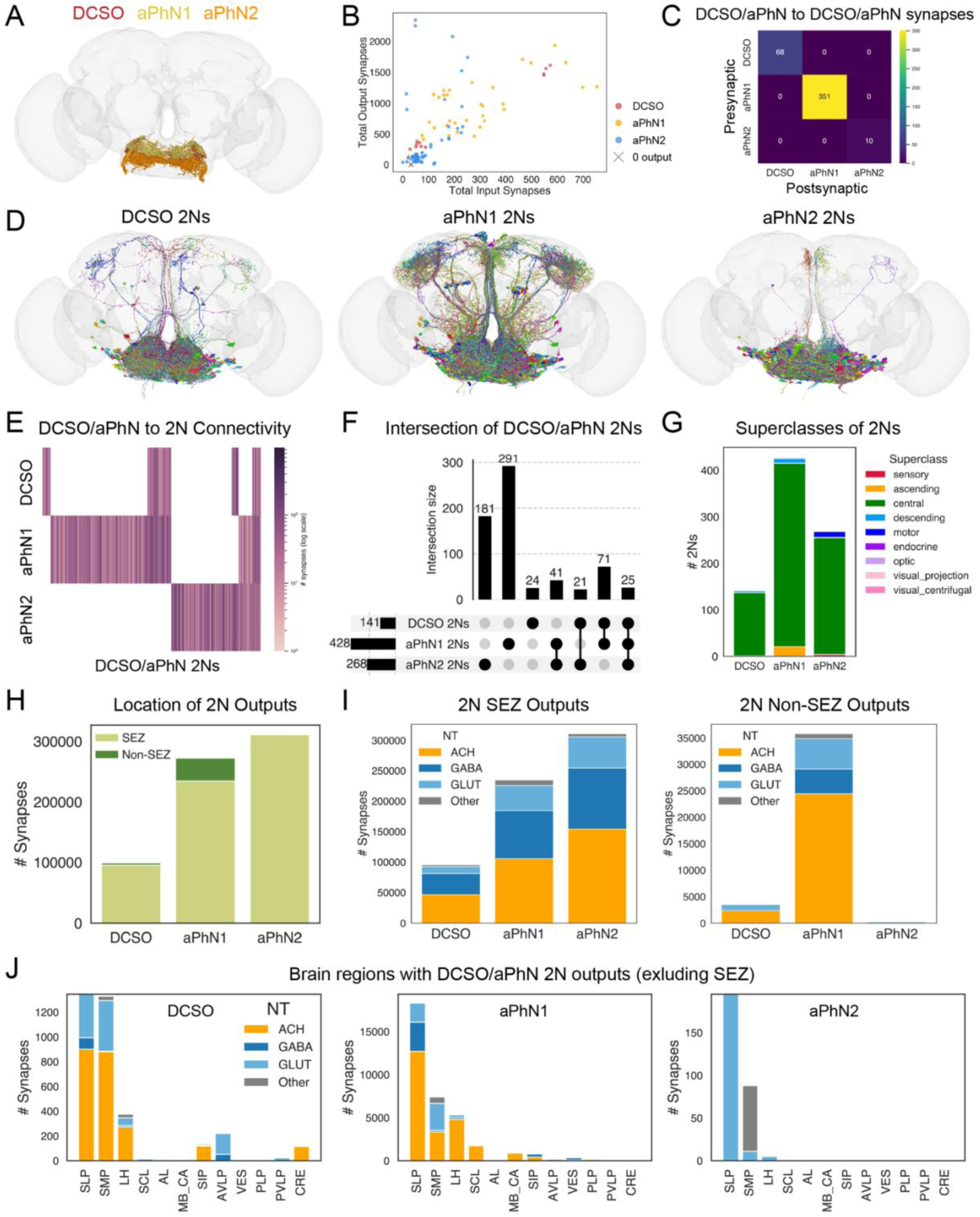
Analyses of the second-order neurons (2Ns) of pharyngeal sensory axons. **(A)** Anterior view of the different groups of pharyngeal sensory neurons: DCSO, aPhN1, and aPhN2. The light gray shading indicates the brain. **(B)** Total output synapses (y-axis) versus input synapses (x-axis) of individual pharyngeal sensory axons. The different groups are indicated by different colors. One axon lacking predicted output synapses is indicated by a cross (X). **(C)** Matrix showing the number of synaptic connections between all pairwise groups of pharyngeal sensory axons. **(D)** Anterior views of the 2Ns of different groups of pharyngeal sensory axons. The light gray shading indicates the brain. (**E-F**) Heatmap (**E**) and UpSet plot (**F**) showing the extent of overlap in 2Ns among different groups of pharyngeal sensory axons. The colors in (**E**) are scaled relative to the number of synapses from each pharyngeal sensory group to each 2N. (**G**) The number of 2Ns belonging to different superclasses. (**H**) The number of 2N output synapses residing within versus outside the SEZ. **(I)** The predicted neurotransmitters (NT) used by the 2N output synapses that reside within the SEZ (left panel) versus outside the SEZ (right panel). ACH, acetylcholine; GABA, gamma-aminobutyric acid; GLUT, glutamate. (**J**) The number of 2N output synapses residing in the indicated non-SEZ brain neuropils for each pharyngeal sensory group. The predicted neurotransmitters (NT) are indicated by different colors.

To examine whether the aPhN1 and aPhN2 axons contain different subtypes, we performed clustering analyses based on the similarity of their postsynaptic partners (Fig. S2), similar to those performed on the StN axons (Fig. 2). These analyses revealed several subtypes of aPhN1 and aPhN2 axons, but they do not differ from each other as distinctly as the StN subtypes do (compare Fig. S2 to Fig. 2). Therefore, we decided not to separately analyze the aPhN1 and aPhN2 subtypes in our subsequent analyses.

The three groups of pharyngeal sensory axons have both distinct and overlapping second-order neurons (2Ns) (Fig. 5D-F). Of the 141 DCSO 2Ns identified, only 24 (17.0%) are unique to DCSO axons; others are shared with aPhN1 and/or aPhN2 axons (Fig. 5E-F).

Interestingly, approximately half of the DCSO 2Ns (71/141) are also aPhN1 2Ns (Fig. 5E-F). This suggests that DCSO and aPhN1 axons, both likely gustatory axons originating from the pharynx, share many postsynaptic partners. On the other hand, larger proportions of 2Ns are unique to aPhN1 (68.0%) and aPhN2 (67.5%) axons (Fig. 5E-F). Nevertheless, common 2Ns are present for any given two groups of pharyngeal sensory axons, and 25 2Ns receive inputs from all three groups (Fig. 5E-F). These results suggest substantial integration of pharyngeal signals at the 2N level.

The DCSO/aPhN 2Ns are primarily central neurons and predicted to use different fast neurotransmitters (Fig. 5G and I). Notably, some motor neurons are directly postsynaptic to aPhN2 axons (Fig. 5G), which will be explored in greater detail in later sections. The majority of output synapses from the DCSO/aPhN 2Ns are located in the SEZ, while non-SEZ output synapses are also present, especially for aPhN1 2Ns (Fig. 5H-I). Outside the SEZ, the superior lateral protocerebrum (SLP) and superior medial protocerebrum (SMP) are the primary targets of DCSO/aPhN 2Ns (Figs. 5D, 5J, and S1). The lateral horn (LH) also receives substantial inputs from DCSO and aPhN1 2Ns (Figs. 5J and S1). These results suggest that pharyngeal sensory signals—particularly the gustatory signals carried by DCSO and aPhN1 axons—are relayed to multiple brain regions outside the SEZ.

The DCSO/aPhN 2Ns synapse onto thousands of 3Ns, covering large proportions of the central brain (Fig. 6A-C). While each group has its unique 3Ns, many 3Ns are shared among two or more groups, suggesting widespread signal integration (Fig. 6B-C). While most of the DCSO/aPhN 3Ns are central neurons, interestingly, a substantial proportion are sensory neurons (Fig. 6D). This suggests that pharyngeal sensory signals may be used to modulate sensory functions. The DCSO/aPhN 3Ns target a wide range of neuropils in addition to the SEZ, using various neurotransmitters (Fig. 6E-G). The SMP, SLP, and LH remain the top targets of DCSO/aPhN 3Ns (Fig. 6G), further implicating these neuropils’ roles in processing pharyngeal sensory signals.

**Figure 6.**
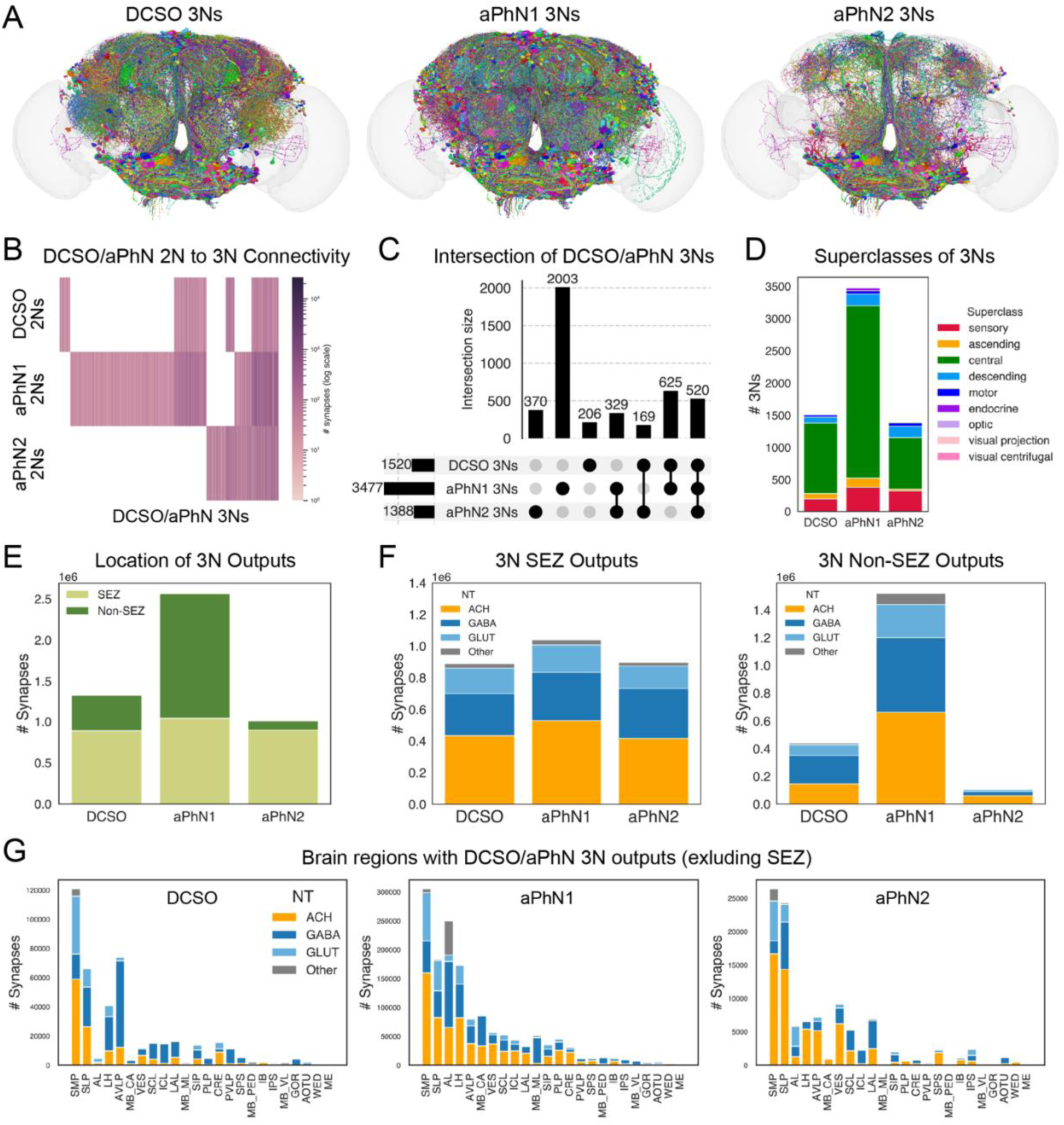
Analyses of the third-order neurons (3Ns) of pharyngeal sensory axons. **(A)** Anterior views of the 3Ns of different groups of pharyngeal sensory axons. The light gray shading indicates the brain. (**B-C**) Heatmap (**B**) and UpSet plot (**C**) showing the extent of overlap in 3Ns among different groups of pharyngeal sensory axons. The colors in (**B**) are scaled relative to the number of synapses from each group of pharyngeal 2Ns to each pharyngeal 3N. (**D**) The number of 3Ns belonging to different superclasses. (**E**) The number of 3N output synapses residing within versus outside the SEZ. (**F**) The predicted neurotransmitters (NT) used by the 3N output synapses that reside within the SEZ (left panel) versus outside the SEZ (right panel). ACH, acetylcholine; GABA, gamma-aminobutyric acid; GLUT, glutamate. (**G**) The number of 3N output synapses residing in the indicated non-SEZ brain neuropils for each pharyngeal sensory group. The predicted neurotransmitters (NT) are indicated by different colors.

### Monosynaptic connections between internal sensory axons and motor or endocrine output neurons

One interesting finding above is that some motor neurons and endocrine cells are directly postsynaptic to gut or pharyngeal sensory neurons (Figs. 3D and 5G). Sensory neurons provide input signals into a circuit, while motor neurons and endocrine cells generate motor and hormonal outputs, respectively. Monosynaptic connections between sensory neurons and motor neurons or endocrine cells indicate the presence of direct input-to-output transformation, which is typically found in simple reflex circuits, such as the knee-jerk reflex ^54^. To further investigate the prevalence of monosynaptic sensory-motor and sensory-endocrine connections, we calculated the minimum number of synaptic hops required for each gut or pharyngeal sensory axon to reach a motor neuron or an endocrine cell (e.g., a value of 1 indicates a monosynaptic connection). For comparison, we performed the same analyses for the four groups of external gustatory receptor neuron (GRNs) annotated in FlyWire: sugar/water, bitter, Ir94e (or low salt), and taste peg. These analyses reveal that the majority of Sets 4-6 StN axons, as well as nearly half of the aPhN2 axons, make monosynaptic connections onto motor neurons (Figs. 7A and S3). On the other hand, the majority of Set 3 StN axons and a few others make monosynaptic connections onto endocrine cells (Figs. 7B and S3). In contrast, none of the external GRNs make monosynaptic connections onto motor neurons or endocrine cells (Figs. 7A-B and S3). These results suggest that a subset of gut and pharyngeal sensory neurons may have more direct influences on motor neurons and endocrine cells than external GRNs.

**Figure 7.**
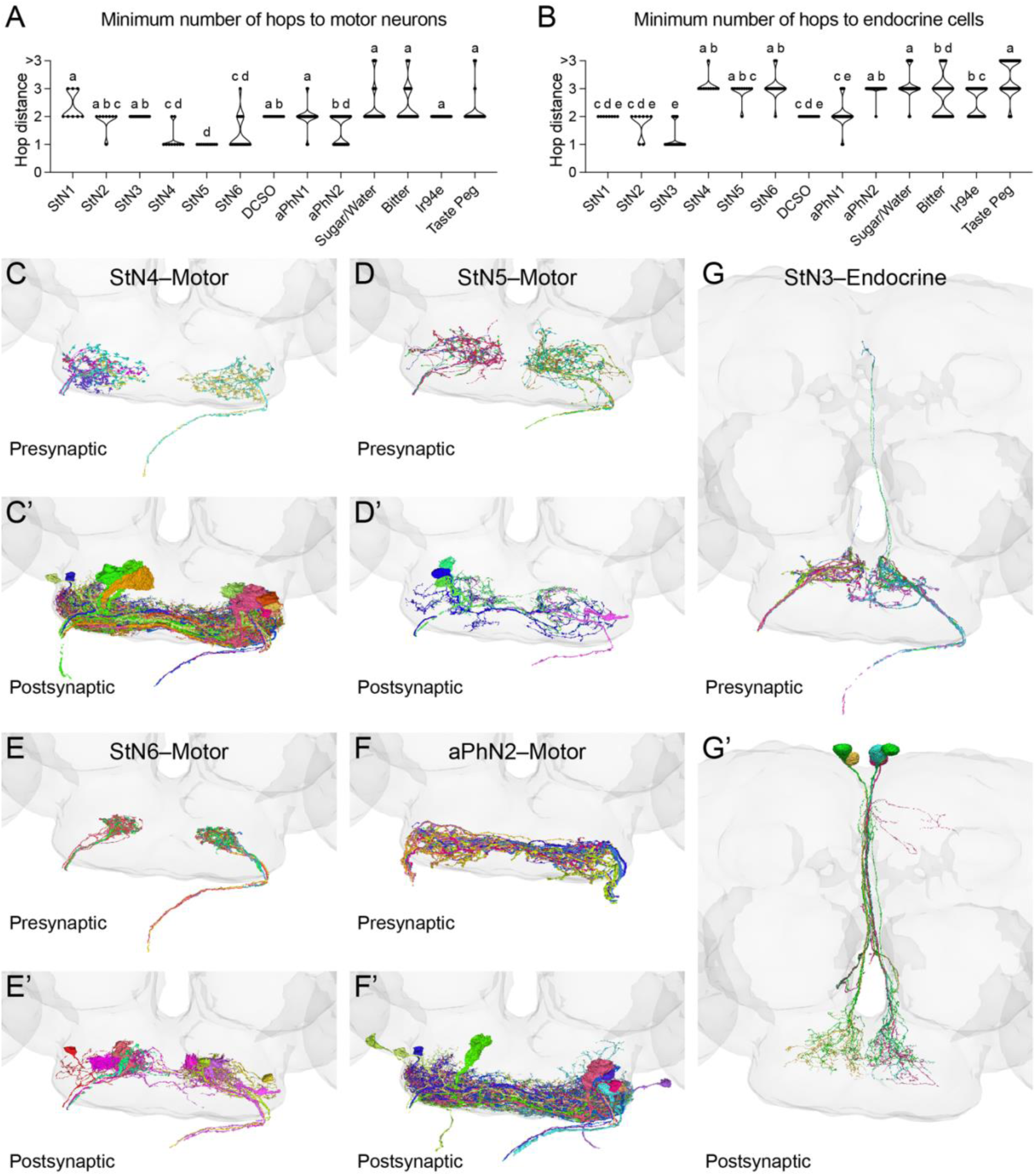
A subset of StN and pharyngeal sensory axons synapses directly onto motor or endocrine neurons. (**A-B**) Violin plots of the minimum number of synaptic hops from the indicated groups of sensory axons to a motor neuron (**A**) or an endocrine cell (**B**). A value of 1 indicates a monosynaptic connection. StN1–6 are different sets of StN axons; DCSO, aPhN1, and aPhN2 are pharyngeal sensory axons; Sugar/Water, Bitter, Ir94e, and Taste Peg are different groups of labellar gustatory receptor neuron (GRNs). See text for details. Different letters indicate significantly different groups (alpha = 0.05), by Kruskal-Wallis test followed by Dunn’s multiple comparisons test. (**C-G’**) Anterior views of the indicated groups of sensory axons (presynaptic) and motor or endocrine neurons (postsynaptic) that form monosynaptic connections. In all panels, the light gray shading indicates the brain.

The Set 4 StN axons form monosynaptic connections with 14 motor neurons, all of which project through the pharyngeal nerves (PhNs) (Fig. 7C-C’). While the identities of these motor neurons are not yet annotated in FlyWire, PhN motor neurons play important roles in food ingestion, including controlling the rhythmic activity of a pharyngeal pump to draw fluid into the esophagus ^27,55–59^. The Set 5 StN axons synapse onto four motor neurons, which also project through the PhNs (Fig. 7D-D’). The identities of these motor neurons are unknown, but they are distinct from the PhN motor neurons postsynaptic to the Set 4 StN axons. The Set 6 StN axons synapse onto nine motor neurons, all projecting through the PhNs (Fig. 7E-E’). Six of these motor neurons are the recently discovered Crop-innervating Enteric Motor (CEM) neurons, which control food entry into the crop, a food storage organ in insects ^34^. This suggests that the Set 6 StN axons may play a role in regulating food storage during ingestion.

Approximately half of the aPhN2 axons make monosynaptic connections onto 11 motor neurons (Fig. 7F-F’). Two of them project through the maxillary-labial nerves (MxLbNs) and are annotated as motor neurons 8 (MN8), which control labella spreading for food ingestion ^58^. The other nine motor neurons project through the PhNs. Most of them have unknown identities, except for one annotated as motor neuron 9 (MN9), which controls the lifting of the rostrum for food reaching ^58,60^. Rostrum lifting and labella spreading are part of the motor programs executed during the feeding initiation phase to assess a food source, but they must also be sustained during the food intake phase ^30^. Our findings above imply that pharyngeal signals from a subset of aPhN2 axons may contribute to sustaining these motor programs, enabling continuous food ingestion. It is also worth noting that the aPhN2 and Set 4 StN axons share some postsynaptic PhN motor neurons (Fig. 7C-C’, F-F’), suggesting signal integration of these two sensory groups at the motor neuron level.

Lastly, the Set 3 StN axons synapse onto five endocrine cells (Fig. 7G-G’), all of which are annotated as median neurosecretory cells that express the *Drosophila* myosuppressin (DMS) peptide. There is a total of six DMS median neurosecretory cells in the adult *Drosophila* brain, which receive sensory inputs through both direct and indirect pathways ^61^. These cells innervate the crop, and DMS modulates crop motility and plays a role in increasing food intake in female flies after mating ^62–66^. Thus, the Set 3 StN axons target the majority of DMS median neurosecretory cells and may play a role in regulating crop physiology and food intake.

## Discussion

In this study, we used the recently completed FAFB connectome ^36–38^ to systemically identify internal sensory axons derived the enteric stomodeal nerve (StN) and different pharyngeal sense organs. These sensory axons play important roles in replaying internal sensory information from the GI tract and pharynx to the brain for feeding regulation, but their anatomy and functions remain incompletely understood. We find that the StN sensory axons contain different subtypes with distinct arborization patterns and output circuits, suggestive of different physiological functions (Figs. 2-4). The pharyngeal sensory axons innervate a more posterior region of the SEZ, with the putative gustatory and mechanosensory axons targeting distinct subregions (Fig. 1). Through analyzing the downstream circuits, we find that the superior medial protocerebrum (SMP) is a top output target of both StN and pharyngeal sensory axons, while the latter group targets additional neuropils (Figs. 3-6). Interestingly, we identify monosynaptic connections from a subset of StN and pharyngeal sensory axons to motor neurons and endocrine cells, suggesting that internal signals from the gut and pharynx may be used to directly control motor and endocrine outputs (Figs. 7 and S3). Together, this work provides a solid foundation for future studies on how diverse internal sensory signals regulate feeding behavior and related endocrine functions.

Future studies are needed to determine whether the StN subtypes identified here have different molecular identities and/or carry different sensory signals. The StN originates from the hypocerebral ganglion (HCG)–corpora cardiaca (CC) region, and carries afferent signals into the brain ^27–30^. The CC contains approximately 13-23 cells expressing the adipokinetic hormone (AKH), which mobilizes energy reserves to maintain circulating sugar levels, similarly to the mammalian glucagon ^33,66–68^. The HCG is less well characterized, estimated to consist of around 15-50 cells by different studies ^32,33,66^. Of these, about 5-6 cells express the mechanosensory receptor Piezo and function to suppress food intake upon gastrointestinal distension ^33^. Another 4-8 HCG cells express the gustatory receptor Gr43a, functioning to detect sugar levels in the gut lumen and direct food entry into the crop ^31–34^. Several neuropeptides/neurotransmitters and their receptors have also been shown to be expressed in the HCG, including sNPF and sNPF-R ^69,70^, DMS and DMS-R1 ^66^, serotonin receptors ^71^, CCHa1-R ^70^, and AstC-R1 ^35^, mediating diverse functions that range from regulating crop motility to sensing sodium salts. This indicates that the HCG contains diverse subtypes that are molecularly and functionally distinct. However, the axonal projections of these HCG subtypes in the central brain are largely uncharacterized. This is probably due to technical challenges, because all the above molecules are also expressed in cells outside the HCG, making it difficult to specifically label the HCG neurons.

Future studies can use intersectional genetic strategies to generate more specific genetic drivers for the HCG subtypes. This may allow better characterization of their axonal patterns and help determine whether the StN subtypes identified here correspond to different molecular subtypes of HCG neurons. Another strategy is to infer the function or identity of the StN subtypes from their connections. For example, the Set 6 StN axons synapse onto four IN1 neurons, which have been shown to receive inputs from HCG neurons expressing the gustatory receptor Gr43a ^34^. This suggests that the Set 6 StN axons are probably the Gr43a-expressing HCG neurons. Interestingly, the Set 6 StN axons directly synapse onto the CEM neurons (Fig. 7), which also receive inputs from the IN1s ^34^, suggesting that the Set 6 StN axons can communicate with the CEM neurons directly or indirectly via the IN1s. Lastly, the Set 6 StN axons form many synapses with each other (Fig. 2C), a feature shared by external gustatory receptor neurons (GRNs), further supporting our speculation that they are gustatory.

The analyses of downstream circuits reveal several brain regions that receive inputs from the putative gut and pharyngeal sensory axons. The SMP is a top output target of both StN and pharyngeal sensory axons (Figs. 3-6), highlighting its importance in processing and integrating internal sensory signals. The SMP is known to receive external gustatory information, as well as internal signals—such as hunger, thirst, and reproductive state—to regulate sugar and water intake ^71–76^. In addition, the SMP is anatomically adjacent to and functionally linked with the pars intercerebralis (PI), a major endocrine structure that secretes multiple peptide hormones involved in regulating feeding, metabolism, and energy homeostasis^77^. A subset of serotonergic neurons, the sugar-SEL projections neurons, anatomically links the SEZ to the SMP ^71^. These neurons are directly postsynaptic to multiple subtypes of StN axons (Sets 4-6) and the aPhN1 and aPhN2 pharyngeal sensory axons (Figs. 3 and 5), suggesting that they may be an important conduit that relays diverse gut and pharyngeal signals to the SMP. The pharyngeal sensory axons, especially the gustatory ones, target additional neuropils, including the SLP and LH (Figs. 5-6). The SLP is a prominent target of taste projection neurons^44,73,78,79^. Our results support this conclusion and further suggest that the SLP is well poised to integrate oral and pharyngeal gustatory information. The LH is best known for its roles in mediating innate behaviors in response to olfactory signals ^80,81^. Our findings that it also receives gustatory inputs from the pharynx imply its potential role in smell-taste integration.

One interesting finding from our studies is that a subset of StN and pharyngeal sensory axons provides monosynaptic inputs onto motor neurons and endocrine cells (Figs. 7 and S3). Monosynaptic connections between sensory and motor neurons are typically found in simple reflex circuits, such as the knee-jerk reflex ^54^. They are thought to underlie rapid and simple signal processing and mediate stereotyped responses. Our findings that several groups of StN and pharyngeal sensory axons form monosynaptic connections with various motor neurons and endocrine cells (Figs. 7 and S3) suggest that internal signals from the gut and pharynx may directly influence feeding-related motor programs and endocrine functions. In contrast, the connections from external gustatory receptor neurons (GRNs) to motor neurons or endocrine cells are always polysynaptic (Figs. 7 and S3). This suggests that the influence of external gustatory signals on motor and endocrine functions may be less direct and subject to integration or modulation, such as by internal states or experience. Similar circuit architectures have also been observed in the larval *Drosophila* feeding connectome ^82–84^. Future studies are invited to better characterize these monosynaptic sensory-motor and sensory-endocrine connections with physiological and behavioral approaches, and determine if they mediate stereotyped, reflex-like responses.

## Resource availability

Requests for further information and resources should be directed to and will be fulfilled by Zepeng Yao.

## Acknowledgments

We thank Meet Zandawala for his help with the cosine similarity clustering analysis, and Philipp Schlegel for discussion on the pharyngeal nerve sensory axons. We thank the labs of Anita Devineni and Gregory Jefferis, and the broader *Drosophila* neuroscience community for sharing connectomics code and annotations. We acknowledge the Princeton FlyWire team and members of Mala Murthy and Sebastian Seung’s labs for the development and maintenance of FlyWire. Members of the Yao lab provided helpful input. This work was supported by startup funds from the University of Florida and a National Institutes of Health (NIH) grant R01DK139131 to Z.Y.

## Author contributions

Z.Y. conceived and supervised the project. D.S.G. performed the majority of the analyses, with input from Z.Y. J.M.Z. and A.J. assisted with the 3D rendering of neurons and figure preparation. Z.Y. and D.S.G. generated the figures, and Z.Y. wrote the manuscript with input from all authors.

## Methods

### Key resources table

| REAGENT OR<br>RESOURCE | SOURCE | IDENTIFIER |
| --- | --- | --- |

| **Retrieved data** |  |  |
| --- | --- | --- |
| FlyWire FAFB connectome (v783 snapshot) | FlyWire Codex | https://codex.flywire.ai/?dataset=fafb |
| FlyWire Codex data export (FAFB) | FlyWire Codex | https://codex.flywire.ai/api/download?dataset=fafb |
| **Software and algorithms** |  |  |
| Python 3.12 | Python Software Foundation | http://www.python.org |
| R (v4.4.2) | R Project | https://www.r-project.org/ |
| Jetbrains Dataspell (Jupyter) | JetBrains | https://www.jetbrains.com/dataspell/ |
| Coconatfly (v0.2.4) | Jefferis Lab | https://github.com/natverse/coconatfly |
| Taste connectome code | Devineni Lab | https://github.com/avdevineni/taste-connectome.git |
| Neuroglancer (3D neuron viewer) | Google | https://github.com/google/neuroglancer |
| GraphPad Prism 10 | GraphPad | https://www.graphpad.com/ |

### Method details

#### Identifying putative gut and pharyngeal sensory axons

We used the FAFB dataset (version 783) in FlyWire ^85^ Codex (https://codex.flywire.ai/?dataset=fafb) to identify sensory neurons in the pharyngeal nerve (PhN) and accessory pharyngeal nerve (aPhN). Specifically, we used search queries “nerve == PhN && super_class == sensory” and “nerve == aPhN && super_class == sensory” to identity the PhN and aPhN sensory axons, respectively. The cell/root IDs and associated metadata, including synapses and annotations, were retrieved from FlyWire Codex. We rendered the identified sensory neurons in 3D within a fly brain mesh using FlyWire’s 3D viewer (https://codex.flywire.ai/app/view_3d?dataset=fafb). We determined their body part origins (StN, DCSO, etc.) and sensory modalities (gustatory or mechanosensory) based on a combination of axon numbers, arborization patterns, nerve entry, and annotations in FlyWire. Details are described in the Results section.

#### Clustering sensory axons based on their output connectivity

We clustered sensory axons based on the similarity of their postsynaptic partners. The cosine similarity method was used to measure similarity, and the Ward D2 implementation of the Ward’s minimum variance hierarchical clustering method was used to perform the hierarchical clustering. At the time of analysis, the default synapse predictions for FlyWire FAFB v783 were generated by Buhmann et al. ^52^; these so-called Buhmann Synapses were used throughout our analyses. In accordance with FlyWire’s suggested filtering convention to remove false positive connections ^37^, only neuronal connections containing a minimum of five synapses are included. Heatmaps and dendrograms were generated using the Coconatfly (v0.2.4) package ^38,86^ in R (v4.4.2) within a DataSpell Jupyter Notebook.

#### Analyzing the second- and third-order downstream neurons

We adapted the method and code used by Walker et al. ^44^ to analyze the second-order neurons (2Ns) and third-order neurons (3Ns) downstream of the gut and pharyngeal sensory axons, with small modifications. Similar to Walker et al. ^44^, we define 2Ns as cells (or root IDs, to be precise) directly postsynaptic to the sensory axons of interest, excluding the first-order sensory axons. Likewise, 3Ns are cells (or root IDs) postsynaptic to 2Ns, excluding 2Ns and the first-order sensory axons. We used a threshold of five synapses for neuronal connections to identify 2Ns and 3Ns. The lists of 2Ns or 3Ns were used to produce heatmaps showing the extent of overlap in 2Ns or 3Ns of the indicated sensory groups. A log-scaled coloring scheme was used to indicate the number of input synapses. These lists were also used to generate UpSet plots to reveal the intersection of 2Ns or 3Ns by neuron count. The 2N and 3N superclasses, synapse locations, and synaptic neurotransmitter predictions were analyzed as previously described ^44^. Following Walker et al. ^44^, we define the SEZ as consisting of the following neuropils: gnathal ganglia (GNG), prow (PRW), saddle (SAD), flange (FLA), and cantle (CAN). All other neuropils are classified as non-SEZ. Neuropil abbreviations are based on Ito et al. ^48^.

#### Analyzing the minimum number of synaptic hops to motor or endocrine neurons

For each root ID in the different sensory groups, we calculated the minimum number of synaptic hops required to reach a motor neuron or an endocrine cell. Only connections containing ≥ 5 synapses were included. A value of 1 therefore indicates a monosynaptic connection. In addition to the internal sensory axons, we analyzed the external GRNs for comparison. The specific search queries used in FlyWire Codex (https://codex.flywire.ai/?dataset=fafb) to retrieve the root IDs for the different groups of external

GRN are as follows:

Sugar/water: nerve == MxLbN && class == gustatory && sub_class == sugar/water Bitter: nerve == MxLbN && class == gustatory && sub_class == bitter

Ir94e (or low salt): nerve == MxLbN && class == gustatory && sub_class == low-salt Taste peg: nerve == MxLbN && class == gustatory && sub_class == taste_peg

The Kruskal-Wallis test, followed by Dunn’s multiple comparisons test, was performed using GraphPad Prism 10 to determine whether the number of hops differs between the sensory groups.

To visualize the downstream circuits of each sensory group, we used linear flow Sankey diagrams to illustrate downstream neurons based on their superclasses, up to three synaptic hops from the sensory neurons. Only connections containing ≥ 5 synapses were included, and feedback connections were not filtered out. Each node and flow represents the total amount of synaptic output connections, with the thickness of the flows scaled relative to the total outflow of their source node.

---

**Figure S1.**
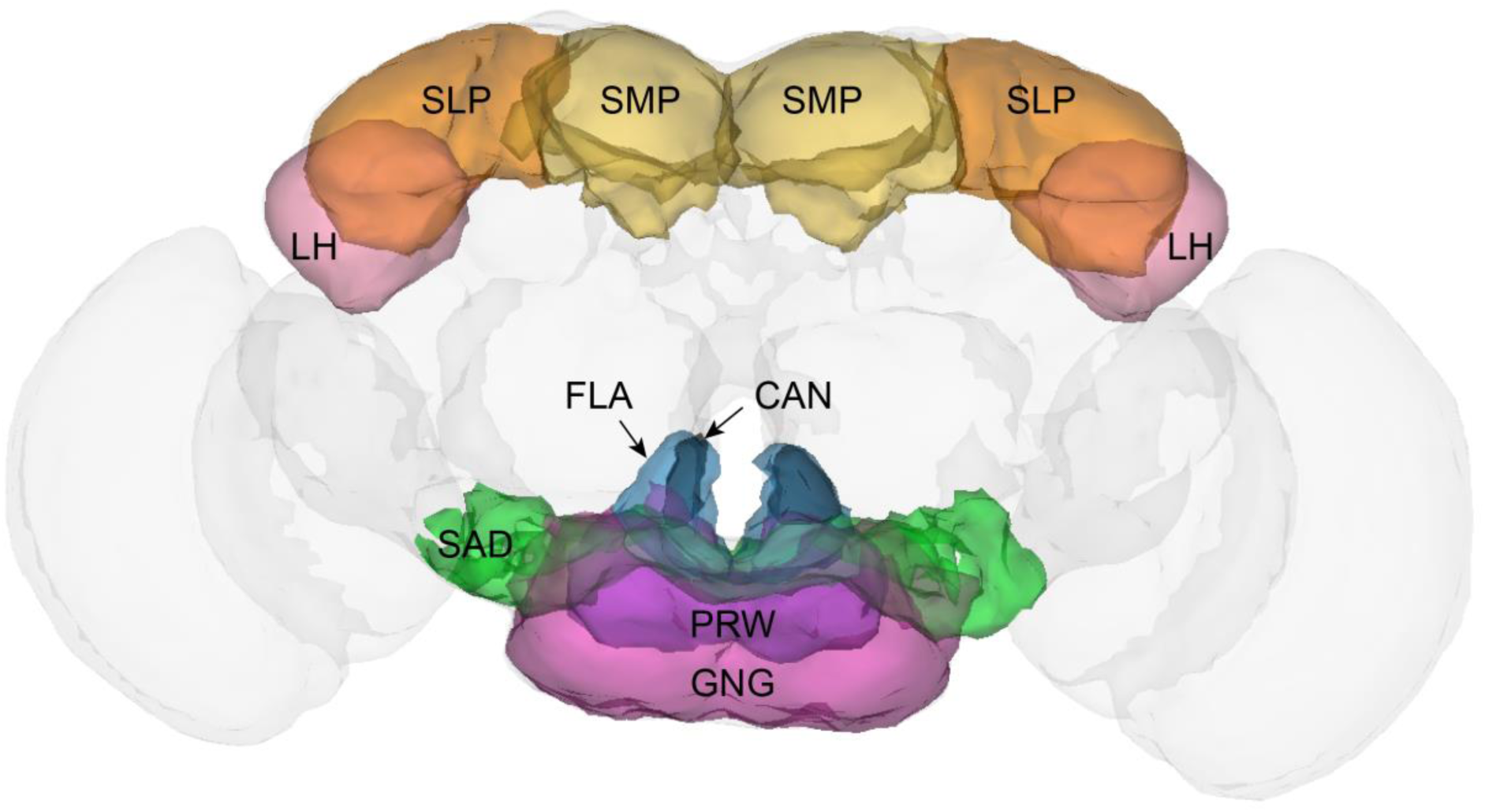
Location of relevant brain neuropils. The light gray shading indicates the fly brain. The relevant neuropils are labeled. SMP, superior medial protocerebrum; SLP, superior lateral protocerebrum; LH, lateral horn. The SEZ consists of GNG (gnathal ganglia), PRW (prow), SAD (saddle), FLA (flange), and CAN (cantle).

**Figure S2.**
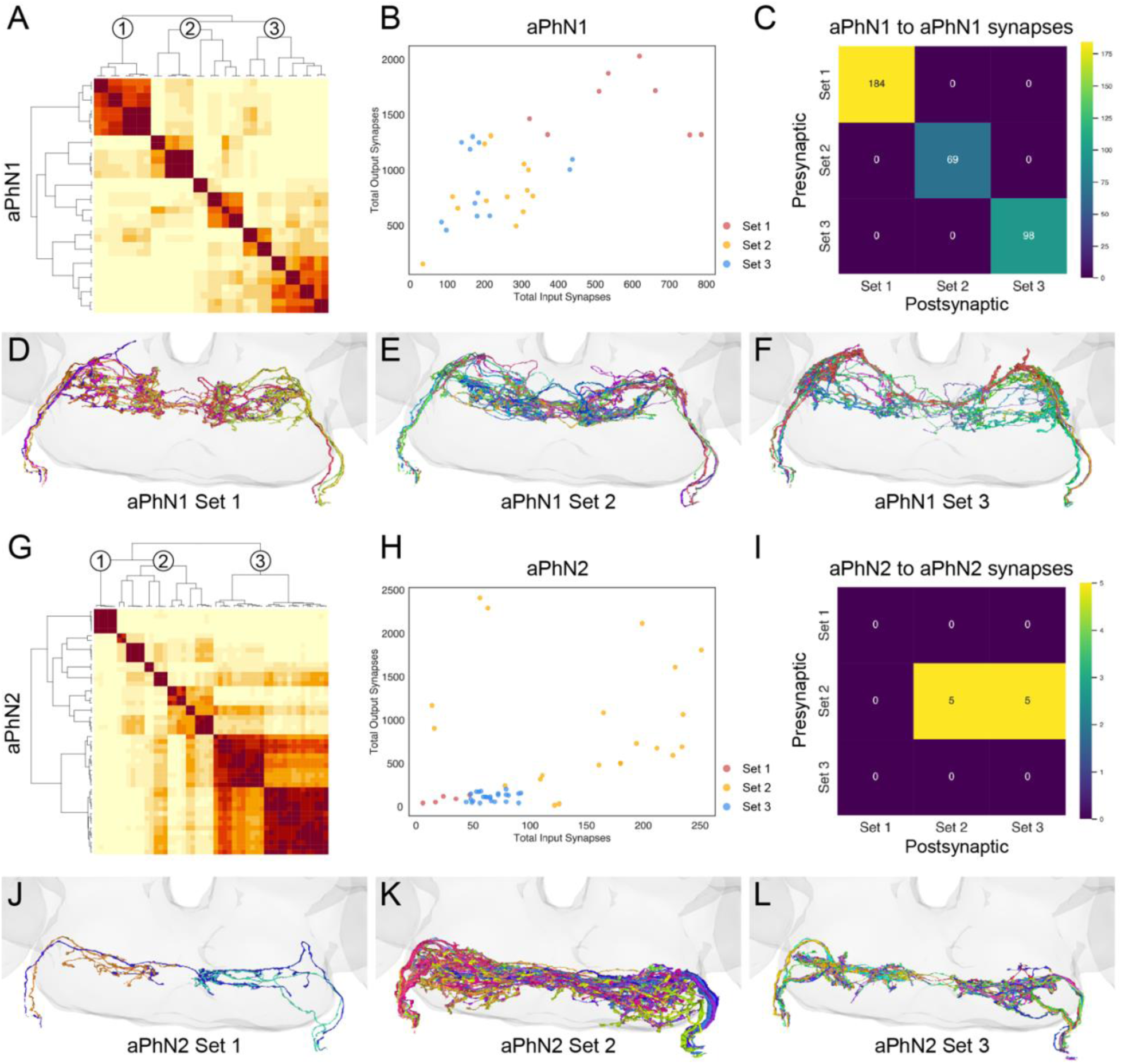
Classification of aPhN1 and aPhN2 sensory axons based on their output synaptic connectivity. (**A, G**) Cosine similarity matrix of aPhN1 axons (**A**) and aPhN2 axons (**G**) based on the similarity of their postsynaptic partners. Darker colors indicate higher similarity between axons. Three sets are identified for each group, as indicated on the dendrograms. (**B, H**) Total output synapses (y-axis) versus input synapses (x-axis) of individual aPhN1 axons (**B**) and aPhN2 axons (**H**). The different sets are indicated by different colors. (**C, I**) Matrices showing the number of synaptic connections between all pairwise sets of aPhN1 axons (**C**) and aPhN2 axons (**I**). (**D-F**) Anterior views of different sets of aPhN1 axons. The light gray shading indicates the brain. (**J-L**) Anterior views of different sets of aPhN2 axons. The light gray shading indicates the brain.

**Figure S3.**
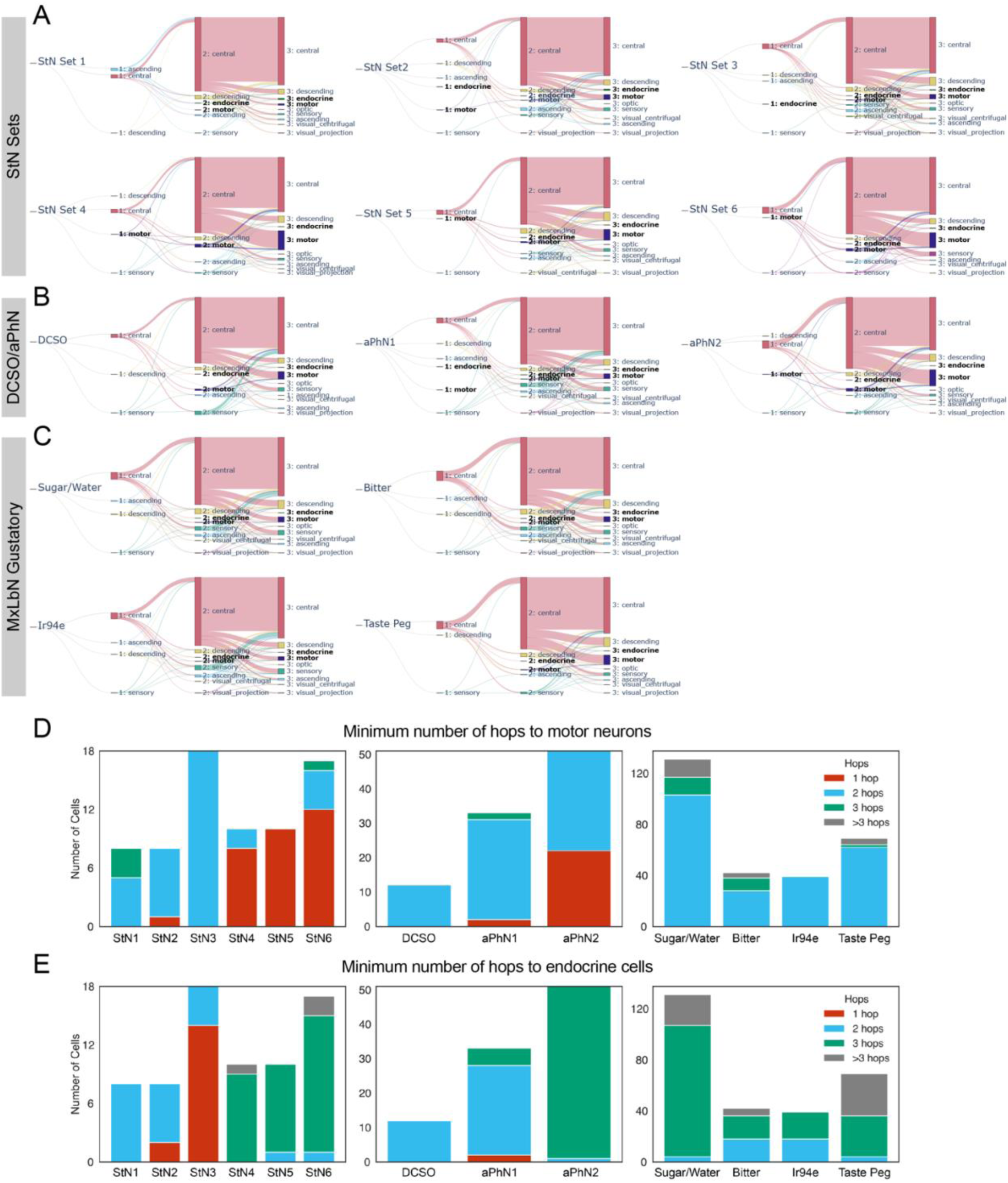
Comparison of downstream circuits between internal and external sensory axons. (**A-C**) Linear flow Sankey diagrams illustrating the downstream neurons grouped by their superclasses, up to three synaptic hops from the indicated groups of sensory neurons. The motor and endocrine superclasses are highlighted in bold. Each node and flow represents the total amount of synaptic output connections, with the thickness of the flows scaled relative to the total outflow of their source node. (**D-E**) Stacked bar charts showing the minimum number of synaptic hops from the indicated groups of sensory axons to a motor neuron (**D**) or an endocrine cell (**E**) (e.g., 1 hop indicates a monosynaptic connection). StN1–6 are different sets of StN axons; DCSO, aPhN1, and aPhN2 are pharyngeal sensory axons; Sugar/Water, Bitter, Ir94e, and Taste Peg are different groups of labellar gustatory receptor neuron (GRNs). See text for details.

## References

1. Yarmolinsky, D.A., Zuker, C.S., and Ryba, N.J.P. (2009). Common Sense about Taste: From Mammals to Insects. Preprint at Elsevier B.V., 10.1016/j.cell.2009.10.001.

2. Chaudhari, N., and Roper, S.D. (2010). The cell biology of taste. Preprint, 10.1083/jcb.201003144.

3. Fontanini, A. (2023). Taste. Preprint at Cell Press, 10.1016/j.cub.2023.01.005.

4. Depoortere, I. (2014). Taste receptors of the gut: Emerging roles in health and disease. Gut 63, 179–190. 10.1136/gutjnl-2013-305112.

5. Steensels, S., and Depoortere, I. (2018). Chemoreceptors in the Gut. Annu Rev Physiol 80, 117–141. 10.1146/annurev-physiol-021317-121332.

6. Kim, M., Heo, G., and Kim, S.Y. (2022). Neural signalling of gut mechanosensation in ingestive and digestive processes. Preprint at Nature Research, 10.1038/s41583-021-00544-7.

7. Mercado-Perez, A., and Beyder, A. (2022). Gut feelings: mechanosensing in the gastrointestinal tract. Preprint at Nature Research, 10.1038/s41575-021-00561-y.

8. Mayer, E.A. (2011). Gut feelings: The emerging biology of gut-”brain communication. Preprint, 10.1038/nrn3071.

9. Alhadeff, A.L., and Yapici, N. (2024). Interoception and gut–brain communication. Preprint at Cell Press, 10.1016/j.cub.2024.10.035.

10. Scott, K. (2018). Gustatory Processing in Drosophila melanogaster. Annu Rev Entomol 63, 15–30. 10.1146/annurev-ento-020117-043331.

11. Chen, Y.C.D., and Dahanukar, A. (2020). Recent advances in the genetic basis of taste detection in Drosophila. Preprint at Springer, 10.1007/s00018-019-03320-0.

12. Shrestha, B., and Lee, Y. (2023). Molecular sensors in the taste system of Drosophila. Preprint at Genetics Society of Korea, 10.1007/s13258-023-01370-0.

13. Park, J.H., and Kwon, J.Y. (2011). Heterogeneous expression of drosophila gustatory receptors in enteroendocrine cells. PLoS One 6. 10.1371/journal.pone.0029022.

14. Ledue, E.E., Chen, Y.C., Jung, A.Y., Dahanukar, A., and Gordon, M.D. (2015). Pharyngeal sense organs drive robust sugar consumption in Drosophila. Nat Commun 6. 10.1038/ncomms7667.

15. Kim, H., Jeong, Y.T., Choi, M.S., Choi, J., Moon, S.J., and Kwon, J.Y. (2017). Involvement of a Gr2a-Expressing Drosophila Pharyngeal Gustatory Receptor Neuron in Regulation of Aversion to High-Salt Foods. Mol Cells 40, 331–338. 10.14348/molcells.2017.0028.

16. Chen, Y.C.D., and Dahanukar, A. (2017). Molecular and Cellular Organization of Taste Neurons in Adult Drosophila Pharynx. Cell Rep 21, 2978–2991. 10.1016/j.celrep.2017.11.041.

17. Chen, Y.C.D., Park, S.J., Joseph, R.M., Ja, W.W., and Dahanukar, A.A. (2019). Combinatorial Pharyngeal Taste Coding for Feeding Avoidance in Adult Drosophila. Cell Rep 29, 961–973.e4. 10.1016/j.celrep.2019.09.036.

18. Chen, Y.C.D., Menon, V., Joseph, R.M., and Dahanukar, A.A. (2021). Control of sugar and amino acid feeding via pharyngeal taste neurons. Journal of Neuroscience 41, 5791–5808. 10.1523/JNEUROSCI.1794-20.2021.

19. Joseph, R.M., Sun, J.S., Tam, E., and Carlson, J.R. (2017). A receptor and neuron that activate a circuit limiting sucrose consumption. Elife 6, 1–25. 10.7554/eLife.24992.

20. Sang, J., Dhakal, S., Shrestha, B., Nath, D.K., Kim, Y., Ganguly, A., Montell, C., and Lee, Y. (2024). A single pair of pharyngeal neurons functions as a commander to reject high salt in Drosophila melanogaster. Elife 12, 93464. 10.7554/eLife.93464.

21. Nayak, S. V., and Singh, R.N. (1983). Sensilla on the tarsal segments and mouthparts of adult Drosophila melanogaster meigen (Diptera: Drosophilidae). Int J Insect Morphol Embryol 12, 273–291. 10.1016/0020-7322(83)90023-5.

22. Gendre, N., Lüer, K., Friche, S., Grillenzoni, N., Ramaekers, A., Technau, G.M., and Stocker, R.F. (2004). Integration of complex larval chemosensory organs into the adult nervous system of *Drosophila*. Development 131, 83–92. 10.1242/dev.00879.

23. Zhang, Y. V., Aikin, T.J., Li, Z., and Montell, C. (2016). The Basis of Food Texture Sensation in Drosophila. Neuron 91, 863–877. 10.1016/j.neuron.2016.07.013.

24. Yang, T., Yuan, Z., Liu, C., Liu, T., and Zhang, W. (2021). A neural circuit integrates pharyngeal sensation to control feeding. Cell Rep 37. 10.1016/j.celrep.2021.109983.

25. Qin, J., Yang, T., Li, K., Liu, T., and Zhang, W. (2024). Pharyngeal mechanosensory neurons control food swallow in Drosophila melanogaster. Elife 12. 10.7554/elife.88614.

26. Yapici, N. (2025). Gut-brain communication in Drosophila melanogaster. Preprint at Elsevier Ltd, 10.1016/j.conb.2025.103096.

27. Miller, A. (1950). The internal anatomy and histology of the imago of Drosophila melanogaster. In The Biology of Drosophila, M. Demerec, ed. (John Wiley & Sons, Inc.), pp. 420–534.

28. Rajashekhar, K.P., and Singh, R.N. (1994). Neuroarchitecture of the tritocerebrum of *Drosophila melanogaster*. Journal of Comparative Neurology 349, 633–645. 10.1002/cne.903490410.

29. Singh, R.N. (1997). Neurobiology of the gustatory systems of Drosophila and some terrestrial insects. Microsc Res Tech 39, 547–563. https://doi.org/10.1002/(SICI)1097-0029(19971215)39:6<547::AID-JEMT7>3.0.CO;2-A.

30. Mahishi, D., and Huetteroth, W. (2019). The prandial process in flies. Curr Opin Insect Sci 36, 157–166. 10.1016/j.cois.2019.09.004.

31. Miyamoto, T., Slone, J., Song, X., and Amrein, H. (2012). A Fructose Receptor Functions as a Nutrient Sensor in the Drosophila Brain. Cell 151, 1113–1125. 10.1016/j.cell.2012.10.024.

32. Miyamoto, T., and Amrein, H. (2014). Diverse roles for the Drosophila fructose sensor Gr43a. Fly (Austin) 8, 19–25. 10.4161/fly.27241.

33. Min, S., Oh, Y., Verma, P., Whitehead, S.C., Yapici, N., Van Vactor, D., Suh, G.S.B., and Liberles, S. (2021). Control of feeding by Piezo-mediated gut mechanosensation in Drosophila. Elife 10, 1–30. 10.7554/eLife.63049.

34. Cui, X., Meiselman, M.R., Thornton, S.N., and Yapici, N. (2024). A gut-brain-gut interoceptive circuit loop gates sugar ingestion in Drosophila. Preprint, 10.1101/2024.09.02.610892.

35. Kim, B., Hwang, G., Yoon, S.E., Kuang, M.C., Wang, J.W., Kim, Y.J., and Suh, G.S.B. (2024). Postprandial sodium sensing by enteric neurons in Drosophila. Preprint at Nature Research, 10.1038/s42255-024-01020-z.

36. Zheng, Z., Lauritzen, J.S., Perlman, E., Robinson, C.G., Nichols, M., Milkie, D., Torrens, O., Price, J., Fisher, C.B., Sharifi, N., et al. (2018). A Complete Electron Microscopy Volume of the Brain of Adult Drosophila melanogaster. Cell 174, 730–743.e22. 10.1016/j.cell.2018.06.019.

37. Dorkenwald, S., Matsliah, A., Sterling, A.R., Schlegel, P., Yu, S.C., McKellar, C.E., Lin, A., Costa, M., Eichler, K., Yin, Y., et al. (2024). Neuronal wiring diagram of an adult brain. Nature 634, 124–138. 10.1038/s41586-024-07558-y.

38. Schlegel, P., Yin, Y., Bates, A.S., Dorkenwald, S., Eichler, K., Brooks, P., Han, D.S., Gkantia, M., dos Santos, M., Munnelly, E.J., et al. (2024). Whole-brain annotation and multi-connectome cell typing of Drosophila. Nature 634, 139–152. 10.1038/s41586-024-07686-5.

39. Bates, A.S., Phelps, J.S., Kim, M., Yang, H.H., Matsliah, A., Ajabi, Z., Perlman, E., Delgado, K.M., Osman, M.A.M., Salmon, C.K., et al. (2025). Distributed control circuits across a brain-and-cord connectome. Preprint, 10.1101/2025.07.31.667571.

40. Berg, S., Beckett, I.R., Costa, M., Schlegel, P., Januszewski, M., Marin, E.C., Nern, A., Preibisch, S., Qiu, W., Takemura, S., et al. (2025). Sexual dimorphism in the complete connectome of the *Drosophila* male central nervous system. Preprint, 10.1101/2025.10.09.680999.

41. Engert, S., Sterne, G.R., Bock, D.D., and Scott, K. (2022). Drosophila gustatory projections are segregated by taste modality and connectivity. Elife 11, 78110. 10.7554/eLife.78110.

42. Shiu, P.K., Sterne, G.R., Engert, S., Dickson, B.J., and Scott, K. (2022). Taste quality and hunger interactions in a feeding sensorimotor circuit. Elife 11. 10.7554/elife.79887.

43. Shiu, P.K., Sterne, G.R., Spiller, N., Franconville, R., Sandoval, A., Zhou, J., Simha, N., Kang, C.H., Yu, S., Kim, J.S., et al. (2024). A Drosophila computational brain model reveals sensorimotor processing. Nature 634, 210–219. 10.1038/s41586-024-07763-9.

44. Walker, S.R., Peña-Garcia, M., and Devineni, A. V. (2025). Connectomic analysis of taste circuits in Drosophila. Sci Rep 15. 10.1038/s41598-025-89088-9.

45. Li, J., Dhaliwal, R., Stanley, M., Junca, P., and Gordon, M.D. (2025). Functional imaging and connectome analyses reveal organizing principles of taste circuits in Drosophila. Current Biology 35, 2391–2405.e4. 10.1016/j.cub.2025.04.035.

46. Tastekin, I., de Haan Vicente, I., Beresford, R.J., Otto, N., Dempsey, G., Waddell, S., and Ribeiro, C. (2025). Connectomics Reveals a Feed-Forward Swallowing Circuit Driving Protein Appetite. Preprint, 10.1101/2025.08.25.671815.

47. Tastekin, I., de Haan Vicente, I., Beresford, R.J., Morris, B.J., Beckett, I., Schlegel, P., Costa, M., Jefferis, G.S.X.E., and Ribeiro, C. (2025). From Sensory Detection to Motor Action: The Comprehensive Drosophila Taste-Feeding Connectome. Preprint, 10.1101/2025.08.25.671814.

48. Ito, K., Shinomiya, K., Ito, M., Armstrong, J.D., Boyan, G., Hartenstein, V., Harzsch, S., Heisenberg, M., Homberg, U., Jenett, A., et al. (2014). A systematic nomenclature for the insect brain. Neuron 81, 755–765. 10.1016/j.neuron.2013.12.017.

49. Wang, Z., Singhvi, A., Kong, P., and Scott, K. (2004). Taste representations in the Drosophila brain. Cell 117, 981–991. 10.1016/j.cell.2004.06.011.

50. Koh, T.W., He, Z., Gorur-Shandilya, S., Menuz, K., Larter, N.K., Stewart, S., and Carlson, J.R. (2014). The Drosophila IR20a Clade of Ionotropic Receptors Are Candidate Taste and Pheromone Receptors. Neuron 83, 850–865. 10.1016/j.neuron.2014.07.012.

51. Kwon, J.Y., Dahanukar, A., Weiss, L.A., and Carlson, J.R. (2014). A map of taste neuron projections in the Drosophila CNS. J Biosci 39, 565–574. 10.1007/s12038-014-9448-6.

52. Buhmann, J., Sheridan, A., Malin-Mayor, C., Schlegel, P., Gerhard, S., Kazimiers, T., Krause, R., Nguyen, T.M., Heinrich, L., Lee, W.-C.A., et al. (2021). Automatic detection of synaptic partners in a whole-brain Drosophila electron microscopy data set. Nat Methods 18, 771–774. 10.1038/s41592-021-01183-7.

53. Eckstein, N., Bates, A.S., Champion, A., Du, M., Yin, Y., Schlegel, P., Lu, A.K.Y., Rymer, T., Finley-May, S., Paterson, T., et al. (2024). Neurotransmitter classification from electron microscopy images at synaptic sites in Drosophila melanogaster. Cell 187, 2574–2594.e23. 10.1016/j.cell.2024.03.016.

54. Reschechtko, S., and Pruszynski, J.A. (2020). Stretch reflexes. Preprint at Cell Press, 10.1016/j.cub.2020.07.092.

55. Rajashekhar, K.P., and Singh, R.N. (1994). Organization of motor neurons innervating the proboscis musculature in Drosophila melanogaster meigen (diptera: Drosophilidae). Int J Insect Morphol Embryol 23, 225–242. 10.1016/0020-7322(94)90020-5.

56. Tissot, M., Gendre, N., and Stocker, R.F. (1998). Drosophila P[Gal4] lines reveal that motor neurons involved in feeding persist through metamorphosis. J Neurobiol 37, 237–250.

57. Manzo, A., Silies, M., Gohl, D.M., and Scott, K. (2012). Motor neurons controlling fluid ingestion in Drosophila. Proc Natl Acad Sci U S A 109, 6307–6312. 10.1073/pnas.1120305109.

58. Schwarz, O., Bohra, A.A., Liu, X., Reichert, H., VijayRaghavan, K., and Pielage, J. (2017). Motor control of Drosophila feeding behavior. Elife 6, 1–32. 10.7554/eLife.19892.

59. McKellar, C.E., Siwanowicz, I., Dickson, B.J., and Simpson, J.H. (2020). Controlling motor neurons of every muscle for fly proboscis reaching. Elife 9, 1–34. 10.7554/eLife.54978.

60. Gordon, M.D., and Scott, K. (2009). Motor control in a Drosophila taste circuit. Neuron 61, 373–384. 10.1016/j.neuron.2008.12.033.

61. McKim, T.H., Gera, J., Gayban, A.J., Reinhard, N., Manoli, G., Hilpert, S., Helfrich-Förster, C., and Zandawala, M. (2024). Synaptic connectome of a neurosecretory network in the Drosophila brain. Preprint, 10.7554/eLife.102684.1.

62. McCormick, J., and Nichols, R. (1993). Spatial and temporal expression identify dromyosuppressin as a brain-gut peptide in Drosophila melanogaster. J Comp Neurol 338, 278–288. 10.1002/cne.903380210.

63. Duttlinger, A., Berry, K., and Nichols, R. (2002). The different effects of three Drosophila melanogaster dFMRFamide-containing peptides on crop contractions suggest these structurally related peptides do not play redundant functions in gut. Peptides (N.Y.) 23, 1953–1957. 10.1016/s0196-9781(02)00179-1.

64. Kaminski, S., Orlowski, E., Berry, K., and Nichols, R. (2002). The effects of three Drosophila melanogaster myotropins on the frequency of foregut contractions differ. J Neurogenet 16, 125–134. 10.1080/01677060213156.

65. Dickerson, M., McCormick, J., Mispelon, M., Paisley, K., and Nichols, R. (2012). Structure-activity and immunochemical data provide evidence of developmental- and tissue-specific myosuppressin signaling. Peptides (N.Y.) 36, 272–279. 10.1016/j.peptides.2012.05.002.

66. Hadjieconomou, D., King, G., Gaspar, P., Mineo, A., Blackie, L., Ameku, T., Studd, C., de Mendoza, A., Diao, F., White, B.H., et al. (2020). Enteric neurons increase maternal food intake during reproduction. Nature 587, 455–459. 10.1038/s41586-020-2866-8.

67. Lee, G., and Park, J.H. (2004). Hemolymph sugar homeostasis and starvation-induced hyperactivity affected by genetic manipulations of the adipokinetic hormone-encoding gene in Drosophila melanogaster. Genetics 167, 311–323. 10.1534/genetics.167.1.311.

68. Hughson, B.N. (2021). The Glucagon-Like Adipokinetic Hormone in Drosophila melanogaster – Biosynthesis and Secretion. Preprint at Frontiers Media S.A., 10.3389/fphys.2021.710652.

69. Oh, Y., Lai, J.S.Y., Mills, H.J., Erdjument-Bromage, H., Giammarinaro, B., Saadipour, K., Wang, J.G., Abu, F., Neubert, T.A., and Suh, G.S.B. (2019). A glucose-sensing neuron pair regulates insulin and glucagon in Drosophila. Nature 574, 559–564. 10.1038/s41586-019-1675-4.

70. Yoshinari, Y., Nishimura, T., Yoshii, T., Kondo, S., Tanimoto, H., Kobayashi, T., Matsuyama, M., and Niwa, R. (2024). A high-protein diet-responsive gut hormone regulates behavioral and metabolic optimization in Drosophila melanogaster. Nature Communications 15. 10.1038/s41467-024-55050-y.

71. Yao, Z., and Scott, K. (2022). Serotonergic neurons translate taste detection into internal nutrient regulation. Neuron 110, 1036–1050.e7. 10.1016/j.neuron.2021.12.028.

72. Jourjine, N., Mullaney, B.C., Mann, K., and Scott, K. (2016). Coupled Sensing of Hunger and Thirst Signals Balances Sugar and Water Consumption. Cell 166, 855–866. 10.1016/j.cell.2016.06.046.

73. Talay, M., Richman, E.B., Snell, N.J., Hartmann, G.G., Fisher, J.D., Sorkaç, A., Santoyo, J.F., Chou-Freed, C., Nair, N., Johnson, M., et al. (2017). Transsynaptic Mapping of Second-Order Taste Neurons in Flies by trans-Tango. Neuron 96, 783–795.e4. 10.1016/j.neuron.2017.10.011.

74. Landayan, D., Wang, B.P., Zhou, J., and Wolf, F.W. (2021). Thirst interneurons that promote water seeking and limit feeding behavior in Drosophila. Elife 10. 10.7554/eLife.66286.

75. González Segarra, A.J., Pontes, G., Jourjine, N., Del Toro, A., and Scott, K. (2023). Hunger- and thirst-sensing neurons modulate a neuroendocrine network to coordinate sugar and water ingestion. Elife 12. 10.7554/eLife.88143.

76. Laturney, M., Sterne, G.R., and Scott, K. (2023). Mating activates neuroendocrine pathways signaling hunger in Drosophila females. Elife 12. 10.7554/eLife.85117.

77. Nässel, D.R., and Zandawala, M. (2020). Hormonal axes in Drosophila: regulation of hormone release and multiplicity of actions. Cell Tissue Res 382, 233–266. 10.1007/s00441-020-03264-z.

78. Kim, H., Kirkhart, C., and Scott, K. (2017). Long-range projection neurons in the taste circuit of Drosophila. Elife 6, e23386. 10.7554/eLife.23386.

79. Deere, J.U., Sarkissian, A.A., Yang, M., Uttley, H.A., Martinez Santana, N., Nguyen, L., Ravi, K., and Devineni, A. V. (2023). Selective integration of diverse taste inputs within a single taste modality. Elife 12. 10.7554/eLife.84856.

80. Schultzhaus, J.N., Saleem, S., Iftikhar, H., and Carney, G.E. (2017). The role of the Drosophila lateral horn in olfactory information processing and behavioral response. J Insect Physiol 98, 29–37. 10.1016/j.jinsphys.2016.11.007.

81. Das Chakraborty, S., and Sachse, S. (2021). Olfactory processing in the lateral horn of Drosophila. Preprint at Springer Science and Business Media Deutschland GmbH, 10.1007/s00441-020-03392-6.

82. Miroschnikow, A., Schlegel, P., Schoofs, A., Hueckesfeld, S., Li, F., Schneider-Mizell, C.M., Fetter, R.D., Truman, J.W., Cardona, A., and Pankratz, M.J. (2018). Convergence of monosynaptic and polysynaptic sensory paths onto common motor outputs in a Drosophila feeding connectome. Elife 7, 1–25. 10.7554/eLife.40247.

83. Miroschnikow, A., Schlegel, P., and Pankratz, M.J. (2020). Making Feeding Decisions in the Drosophila Nervous System. Preprint at Cell Press, 10.1016/j.cub.2020.06.036.

84. Schoofs, A., Miroschnikow, A., Schlegel, P., Zinke, I., Schneider-Mizell, C.M., Cardona, A., and Pankratz, M.J. (2024). Serotonergic modulation of swallowing in a complete fly vagus nerve connectome. Current Biology 34, 4495–4512.e6. 10.1016/j.cub.2024.08.025.

85. Dorkenwald, S., McKellar, C.E., Macrina, T., Kemnitz, N., Lee, K., Lu, R., Wu, J., Popovych, S., Mitchell, E., Nehoran, B., et al. (2022). FlyWire: online community for whole-brain connectomics. Nat Methods 19, 119–128. 10.1038/s41592-021-01330-0.

86. Jefferis G (2025). coconatfly: Comparative Connectomics of Public and In Progress Drosophila Datasets. R package version 0.2.4.9000, https://github.com/natverse/coconatfly. Preprint.

